# Comprehensive analysis of prime editing outcomes in human embryonic stem cells

**DOI:** 10.1101/2021.04.12.439533

**Authors:** Omer Habib, Gizem Habib, Gue-Ho Hwang, Sangsu Bae

## Abstract

Prime editing is a versatile and precise genome editing technique that can directly copy desired genetic modifications into target DNA sites without the need for donor DNA. This technique holds great promise for the analysis of gene function, disease modeling, and the correction of pathogenic mutations in clinically relevant cells such as human pluripotent stem cells (hPSCs). Here we comprehensively tested prime editing in hPSCs by generating a doxycycline-inducible prime editing platform. Prime editing successfully induced all types of nucleotide substitutions and small insertions and deletions, similar to observations in other human cell types. Moreover, we compared prime editing and base editing for correcting a disease-related mutation in induced pluripotent stem cells derived form a patient with α 1-antitrypsin (A1AT) deficiency. Finally, whole-genome sequencing showed that, unlike the cytidine deaminase domain of cytosine base editors, the reverse transcriptase domain of a prime editor does not lead to guide RNA-independent off-target mutations in the genome. Our results demonstrate that prime editing in hPSCs has great potential for complementing previously developed CRISPR genome editing tools.

## Introduction

Human pluripotent stem cells (hPSCs), including embryonic stem cells (ESCs) and induced pluripotent stem cells (iPSCs), have the capacity to differentiate into cells derived from the three embryonic germ layers. The ability to engineer the genome of hPSCs has led to the generation of powerful *in vitro* systems in the fields of developmental biology, disease modeling, drug discovery, and stem cell-based therapeutics^1-4^.

Recently, the CRISPR-Cas9 system has been developed as a major tool for genome editing. This system relies on customizable RNAs that guide Cas9 nucleases to target DNA sequences, resulting in the generation of a double strand break (DSB)^5-8^; the response of the cell’s repair systems, such as nonhomologous end-joining (NHEJ) or homology-directed repair (HDR), leads to targeted genome editing. However, it has been reported that even a single DSB can induce chromosomal rearrangements or large deletions^9^, and, especially in hPSCs, can lead to the activation of the p53 pathway, resulting in apoptosis^10^. Moreover, the efficiency of genome editing via CRISPR-Cas9-induced DSBs is generally low in hPSCs^8, 11^. Alternatively, DNA base editors (BEs), including cytosine base editors (CBEs) and adenine base editors (ABEs) can generate single-nucleotide substitutions without introducing a DSB^12^. But these tools have potential limitations, such as a restricted editing window, bystander nucleotide editing, and applications confined to transition mutations (C•G to T•A conversions for CBEs and A•T to G•C conversions for ABEs)^13^. Furthermore, several studies have revealed genome-wide CBE-mediated off-target DNA editing, transcriptome-wide CBE- and ABE-mediated off-target RNA editing, and unwanted, incorrect editing such as ABE-mediated cytosine deamination^14-18^.

Prime editors (PEs) are newly developed genome editing tools that can introduce all types of base-to-base conversions (i.e., transition and transversion mutations) as well as small insertions and deletions (indels) without inducing DSBs^19^. PE2 consists of a Cas9 nickase (nCas9) containing a H840A mutation and an engineered reverse transcriptase. PE2 employs a prime editing guide RNA (pegRNA) to find its target site and installs the desired edit at the site. The pegRNA, an engineered form of standard single-guide RNA (sgRNA), has an extension sequence at the 3’ end that includes a primer binding site (PBS) that primes the RT reaction and an RT template that encodes the desired edit. When the PE-pegRNA complex binds to the target DNA, the nCas9 domain nicks the strand containing the protospacer adjacent motif (PAM) and generates a 3’ end at the target site. Next, the liberated 3’ end hybridizes to the PBS of the pegRNA to prime the RT reaction, after which the RT incorporates the desired sequences into the target using the pegRNA as a template. Annealing of the newly generated 3’ flap with the non-edited strand exposes a 5’ flap that is subsequently removed by the DNA repair machinery (Supplementary Fig. 1). Introduction of a second nick into the opposite strand with an additional sgRNA (a nicking sgRNA) can further improve prime editing efficiency; systems that incorporate a nicking sgRNA are designated as PE3. Thus, both PE2 and PE3 are capable of copying information directly from the pegRNA into the target DNA sequence. Recently, a study harnessed PEs to edit a reporter gene in hiPSCs^20^, but PEs have not been extensively tested or characterized in hPSCs.

Here, we constructed a drug-inducible, PE2-expressing hESC line. We demonstrated the value of this PE platform for introducing small indels and all types of nucleotide substitutions at target sites by testing 66 pegRNAs. In addition, we compared PEs with BEs for the correction of a disease-causing mutation in patient-derived iPSCs. Finally, through whole genome sequencing (WGS), we revealed that prolonged expression of PE2 in hPSCs does not result in the generation of unintended mutations in the genome.

## Results

### Construction of a doxycycline-inducible, PE2-expressing hPSC line named H9-iPE2

To systematically investigate the editing features of PEs in hPSCs, we constructed an hPSC line in which PE2 expression can be induced by doxycycline (dox). First, we prepared a construct that contains the PE2 sequence under the control of an inducible TetO promoter and a neomycin selection cassette (Fig. 1a). The PE2 construct was integrated into the first intron of the *PPP1R12C* gene (also known as the AAVS1 site) in human H9 ESCs by using transcription activator-like effector nuclease (TALEN)-mediated gene targeting. After neomycin selection, individual colonies were picked and expanded for genotyping. We examined whether the PE2 construct was properly integrated into the target site using PCR with specific primer sets spanning the junctions between the endogenous sequences and the integrated donor DNA (Fig. 1b) and confirmed the junctions by Sanger sequencing (Supplementary Fig. 2). For a selected cell line named H9-iPE2, we found using immunostaining that PE2 was specifically expressed in the presence of dox (Fig. 1c). To examine the editing capacity of the H9-iPE2 cells, we first determined prime editing efficiencies with three HEK3-targeting pegRNAs (Supplementary datasheet), which were previously used in HEK293T cells^19^. Due to its higher editing efficiency compared with that of PE2, we used the PE3 system throughout the study. The H9-iPE2 cells were electroporated with each pegRNA and a nicking sgRNA (targeting a site +63 nt away from the 5’ end of the pegRNA-induced nick) and treated with dox for 3 days in a 48-well plate. Then, high-throughput sequencing was used to quantify the editing efficiency in bulk populations. The results revealed that PE3 introduced the intended mutations with editing efficiencies between 4.3–24.8%, supporting the editing capacity of the H9-iPE2 cells (Fig. 1d). We also observed that PE3 generated unwanted indels at a frequency of 2.7 to 8.1%.

**Fig. 1.**
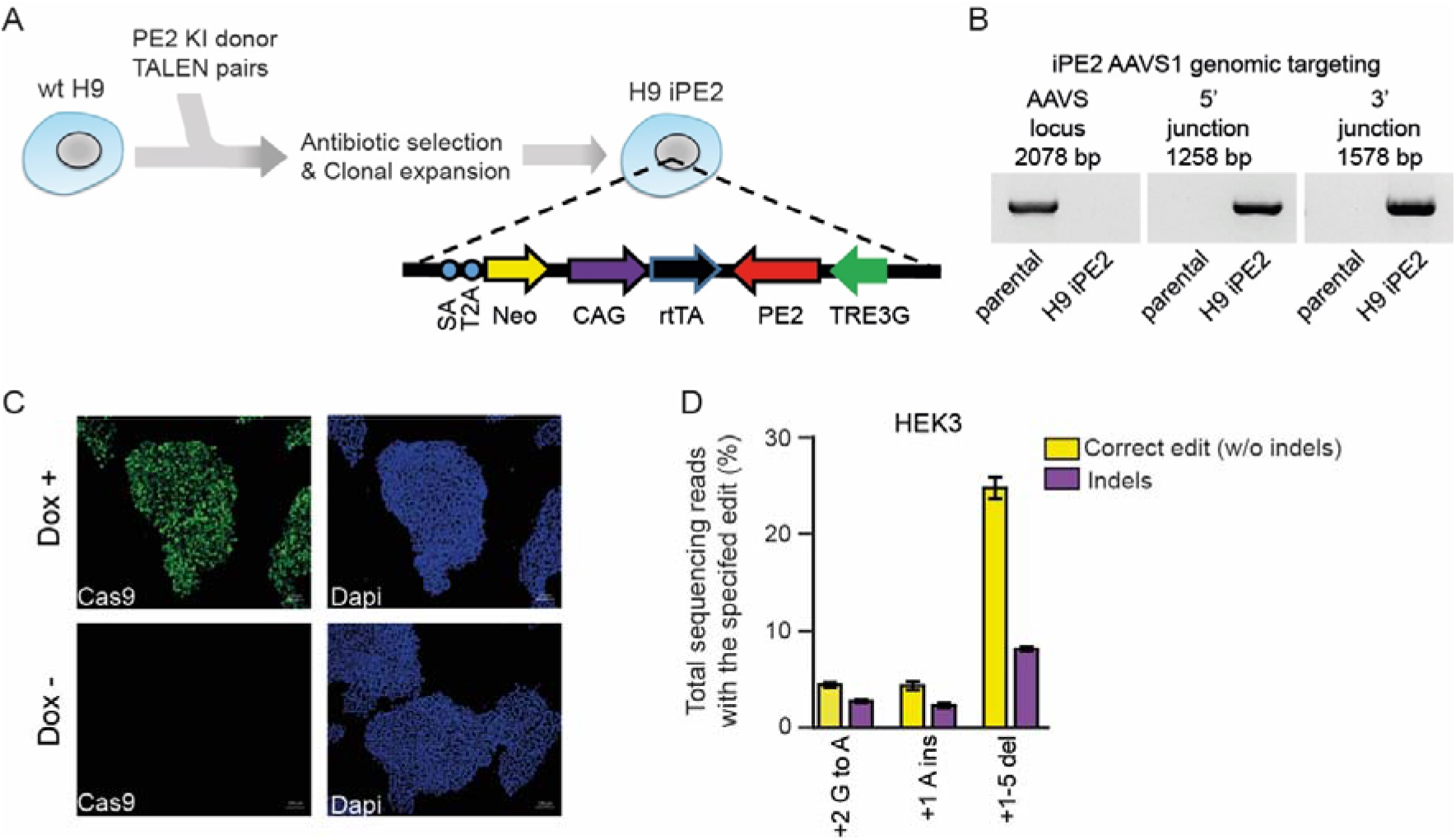
Generation and characterization of H9-iPE2. **a** Schematic diagram of the strategy for TALEN-mediated targeting of the AAVS1 locus to generate H9-iPE2 cells, in which PE2 expression is induced by dox. The AAVS1 donor vector contains a cassette in which PE2 expression is under the control of the dox-inducible TRE3G promoter. SA, splice acceptor; 2A, self-cleaving 2A peptide; Neo, neomycin resistance gene; rtTA, dox-controlled reverse transcriptional activator; CAG, cytomegalovirus early enhancer/chicken β actin promoter. **b** PCR-based confirmation that the iPE2 construct was targeted to the AAVS1 locus. A primer pair that flanks the AAVS1 knock-in site only amplifies a product in wild-type cells; lack of a product indicates that the H9-iPE2 clone used in this study is homozygous (left panel). Junction PCR confirmed on-target integration of the cassette into the AAVS1 locus (center and right panels). **c** Immunostaining of H9-iPE2 cells before and after 48 h of dox treatment with an anti-Cas9 antibody (green). Nuclei were stained with DAPI (blue). **d** Editing efficiency in H9-iPE2 cells. H9-iPE2 cells maintained in the presence of dox were electroporated with plasmids encoding a pegRNA and a nicking sgRNA. The editing efficiency is indicated as the percentage of total sequencing reads that contain the intended edit and do not contain indels. Mean ± s.d. of n = 3 independent biological replicates.

### Precise PE-mediated editing at endogenous loci in hPSCs

We next investigated various features of PE3-mediated editing in H9-iPE2 cells. It is known that the PE-mediated editing efficiency is dependent on the lengths of both the PBS and RT template in the pegRNA^19^. To examine this point in H9-iPE2 cells, we selected target sites at two different loci, HEK3 and RNF2, and tested pegRNAs with varying PBS lengths together with corresponding nicking sgRNAs. In a previous study using HEK293T cells, the Liu group reported that efficient prime editing at the RNF2 site, which has a relatively low G/C content (35%), required a longer PBS sequence than did efficient prime editing at the HEK3 site, which has a G/C content of 71%. Similarly, we found that efficient prime editing at the RNF2 site required a longer PBS sequence than at the HEK3 site (Figs. 2a and 2b), suggesting that the optimum PBS length is sequence dependent and might be conserved among cell types (Supplementary Fig. 3). In addition, similar to results from the previous study in HEK293T cells, we found that PEs can introduce both transition and transversion substitutions across the +1 to +30 positions of the HEK3 target site with varying efficiencies in hPSCs (Figs. 2c and 2d). We also tested the ability of PE3 to introduce small indels (1 to 3 bps) at the HEK3 and RNF2 sites and found that PE3 successfully generated the intended edits (Figs. 2e and 2f). Finally, we demonstrated that large deletions (up to 80 bps) and insertions (up to 42 bps) were precisely installed by PE3 (Figs. 2g and 2h). From these results, we concluded that PE3 can be a practical tool for inducing a broad range of mutations in hPSCs.

**Fig. 2.**
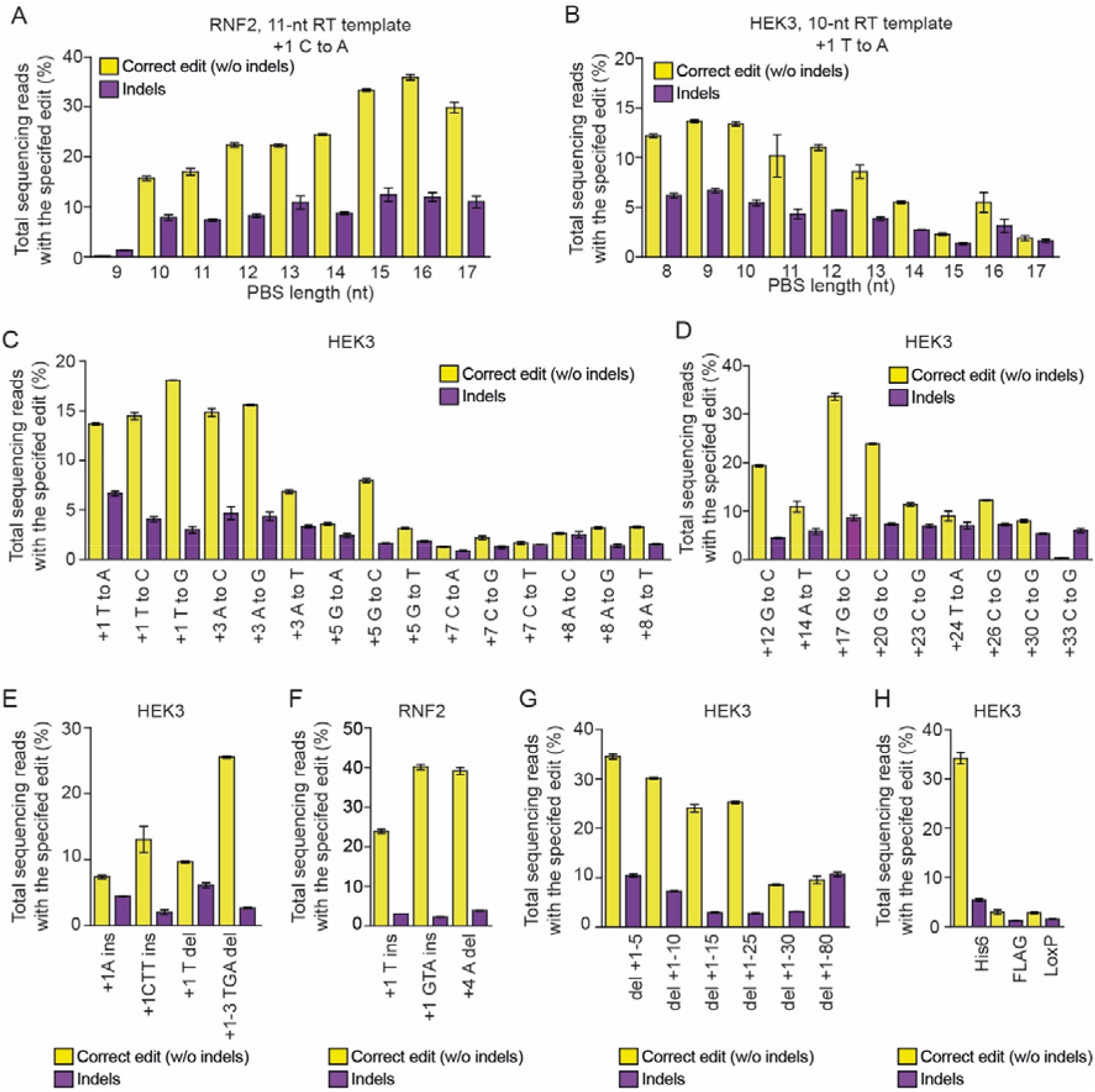
Prime editing of genomic DNA in H9-iPE2 cells by PE3. **a-b** PE3 editing efficiencies at HEK3 and RNF2 genomic sites with pegRNAs containing varying PBS lengths. **c** Efficiencies of all 12 types of transition and transversion edits at the indicated positions in the HEK3 site. **d** Efficiencies of long distance edits at the HEK3 site using a 34-nt RT template. **e-f** Editing efficiencies for the generation of intended indels at the HEK3 and RNF2 genomic sites. Yellow, desired indel; purple, undesired indel. **g** Editing efficiencies for the generation of targeted deletions of 5–80 bps at the HEK3 site. **h** Editing efficiencies for the generation of targeted insertions of a His6 tag, Flag epitope tag, and loxP site at the HEK3 site. The editing efficiency is indicated as the percentage of total sequencing reads that contain the intended edit and do not contain unintended indels. Mean ± s.d. of n = 3 independent biological replicates.

### Double nicking strategy of PE3 is not the primary reason for unwanted indel generation

We next sought to understand the reason why indel mutations are generated by the PE3 system. To investigate which factor has a dominant role for generating such indel patterns, we examined five different scenarios: i) a single DSB induced by wild-type (wt) Cas9 and a sgRNA, ii) a single nick induced by nCas9-RT and a sgRNA, iii) double DSBs induced by wt Cas9 and two sgRNAs, iv) double nicks induced by nCas9-RT and two sgRNAs, and v) double nicks induced by nCas9-RT, one pegRNA, and one sgRNA (i.e., the PE3 system) (Fig. 3a). We tested all of these possibilities at both the HEK3 and RNF2 sites. The distances between the first and second nicks were 63 and 41 nt for the HEK3 and RNF2 sites, respectively. To maintain consistency with the PE3 system, we additionally established an H9 line in which wt Cas9 was expressed in a dox-inducible manner, named H9-iCas9 (Supplementary Fig. 4). We also set the ratio of the first sgRNA to the second sgRNA at 3:1.

**Fig. 3.**
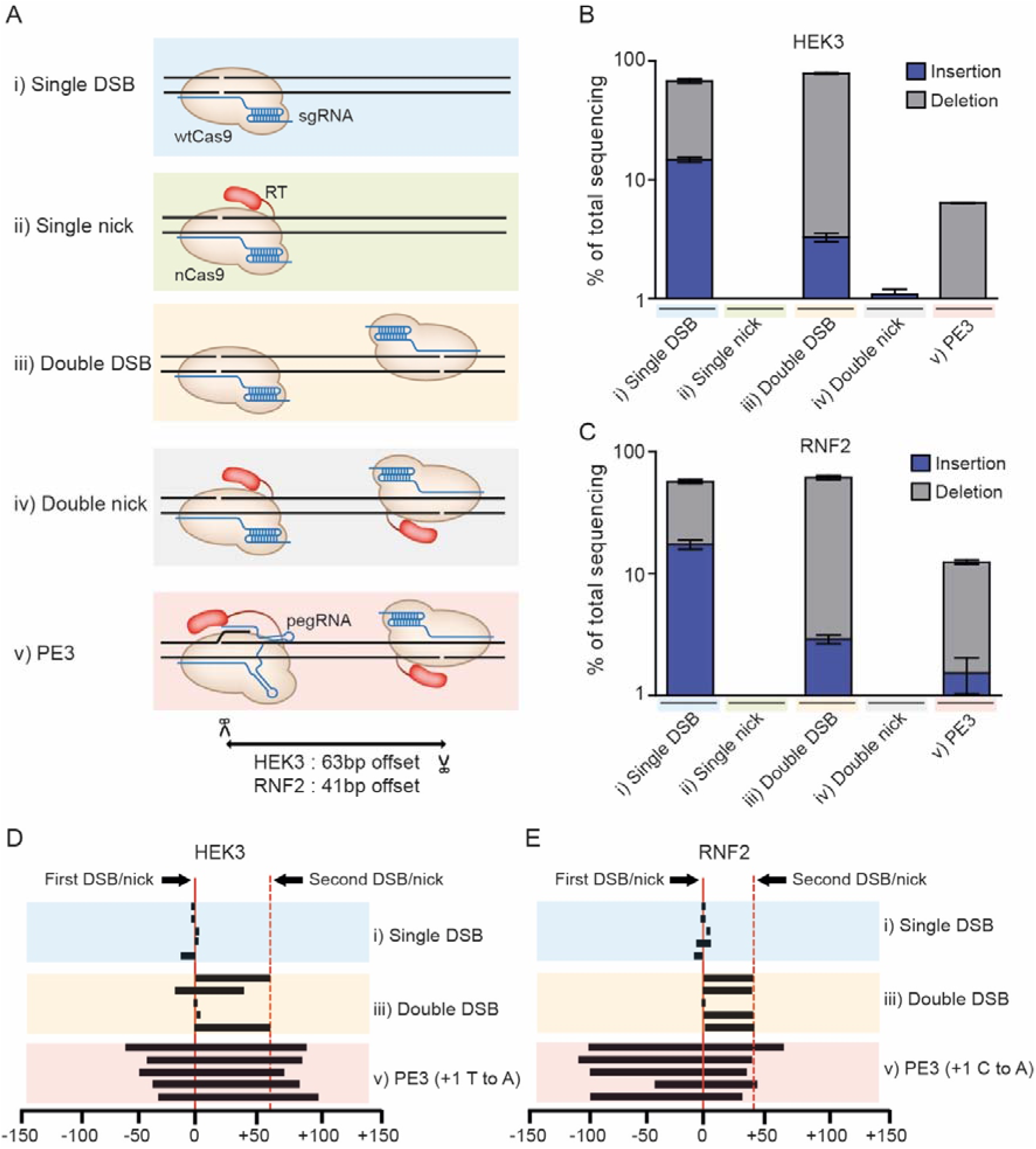
Comparison of indel frequencies and endpoints after different DNA cleavage and nicking scenarios. (**A**) Schematic of enzymes and guide RNAs tested to compare indel frequencies. **b-c** Indel frequencies generated by the experimental set ups diagrammed in A at the HEK3 and and RNF2 genomic sites. **d-e** Deletion patterns obtained after single cuts, double cuts, and PE3-mediated base substitutions at the HEK3 and RNF2 sites. Each line designates the extent of a deletion. The five most frequent types of deletions are shown for each case.

We first assessed indel frequencies after the introduction of identical sgRNAs into H9-iCas9 and H9-iPE2 cells. High-throughput sequencing data showed that the resulting single DSBs (scenario i) led to high frequencies of indels (68% and 79%) at the HEK3 and RNF2 sites, respectively, whereas no significant indel frequencies were observed after the introduction of single nicks (ii) (Figs. 3b and 3c). We also electroporated a pair of sgRNAs to examine the effects of double DSBs or nicks. The sequencing data showed that double DSBs (iii) resulted in high indel frequencies, but that double nicks (iv) did not (indel frequencies of <2%), suggesting that the generation of double nicks in and of itself is not a major reason why indels are generated and that the RT enzyme in PE3 cannot generate indel mutations alone. However, when double nicks were produced following the introduction of a pegRNA and a sgRNA (v), the frequency of indels substantially increased (up to 12%), suggesting that the combination of the RT enzyme and a pegRNA could cause indel mutations. We further investigated the indel patterns generated following a single DSB (i), double DSBs (iii), and the double nicks introduced by PE3 (v). Five recurrent deletion patterns were obtained in each situation for the comparison. PE3 generated long deletions that included the region upstream of the first DSB site, whereas a single DSB led to the generation of small deletions near the DSB site and double DSBs typically induced precise deletion of the region between the two DSB sites (Figs. 3d and 3e). These results also supported our conclusion that unintended indels are not directly caused by double nicking per se but by the combinatory activity of the RT and the pegRNA.

### Comparison of prime editing and base editing in hPSCs

We next compared the activity of PEs and BEs at two genomic sites (HEK3 and FANCF) in hPSCs. In this experiment, transient BE overexpression was achieved by electroporating BE-encoding plasmids into parental H9 cells, whereas the H9-iPE2 cells were used for assessing prime editing. We first measured the activity of BE4max, an optimized CBE variant, at both the HEK3 and FANCF sites. The high-throughput sequencing data showed that several cytosines within the editing window were edited at a frequency of up to 23% at the HEK3 site (Fig. 4a) and 26% at the FANCF site (Fig. 4b), indicating CBE bystander effects. In contrast, for the PE experiment, we tested individual pegRNAs designed to convert each cytosine at both the HEK3 and FANCF sites. The high-throughput sequencing data showed that each pegRNA led to precise editing of the targeted cytosine, although the conversion rates of the PEs were typically lower than that of the CBEs (Figs. 4a and 4b). We also tested ABEs (ABEmax) at the same target sites, and found that they exhibited similar tendencies as the CBEs (Figs. 4c and 4d). In these experiments, we found that both CBEs and ABEs generally showed higher activities than PEs, although PE3 has an advantage in that the PE2 system is expressed in virtually all cells. However, we also found that bystander nucleotide conversion by BEs was inevitable when the target contained multiple cytosines or adenines in the editing window, whereas PEs precisely converted target nucleotides. We also measured indel frequencies in all experiments and found that PE3s were associated with apparent indels (at frequencies ranging from 2% to 15%), whereas CBEs induced indels at a frequency of 2% and ABEs led to an insignificant indel frequency (Figs. 4e-4h). Taken together, our results indicate that because of their high activity, BEs are preferred over PEs for targets that lack bystander nucleotides, such as the FANCF site for ABE. However, PE-induced base conversion is generally more precise, although indel generation can occur in the case of the PE3 system.

**Fig. 4.**
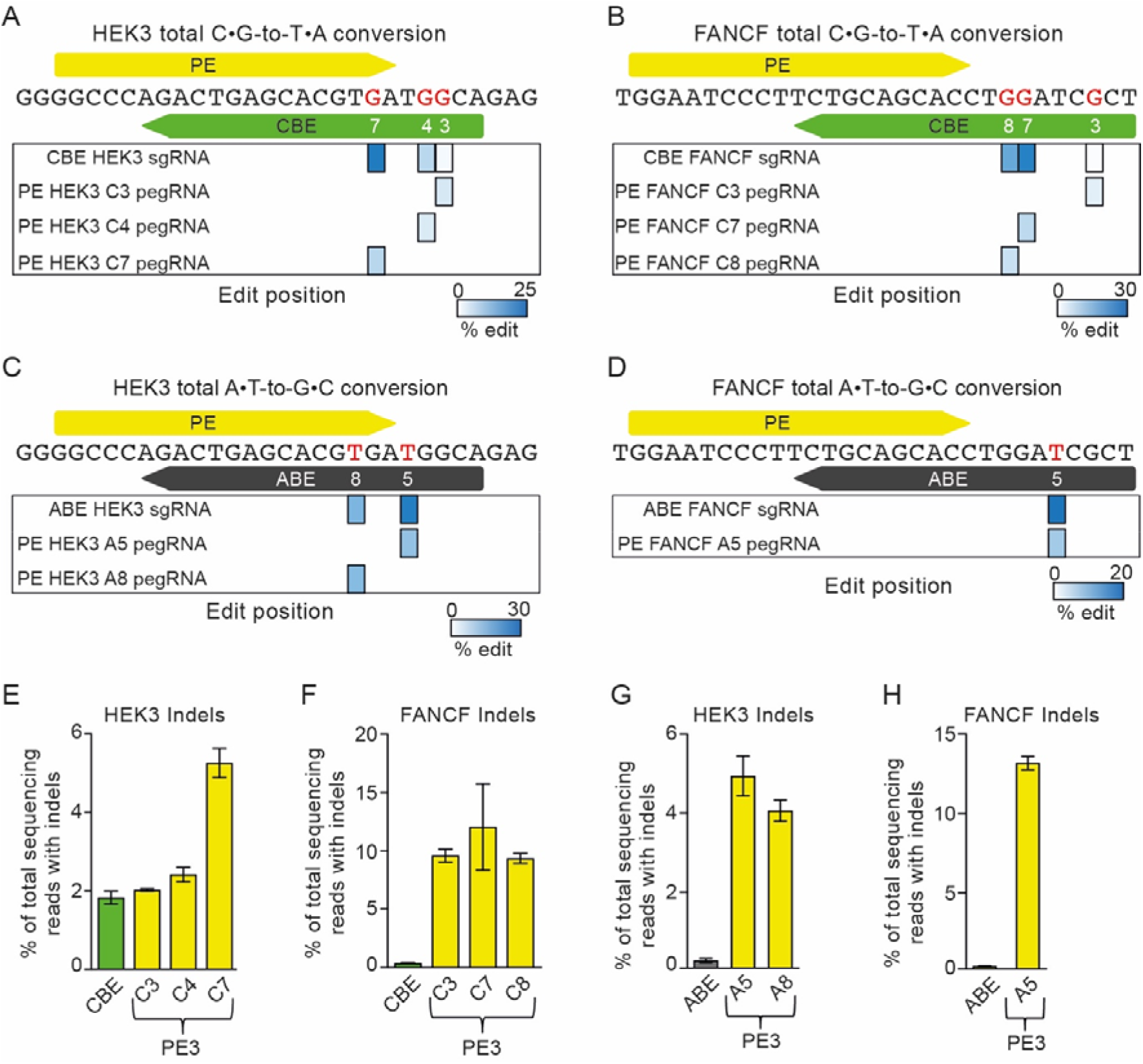
Comparison of prime editing and base editing outcomes at the same target sites. **a-b** CBE- and PE-generated frequencies of C•G-to-T•A edits at target nucleotides, highlighted in red, in the endogenous HEK3 and FANCF sites. Base positions are numbered relative to the PAM of the CBE sgRNA. The PAM nucleotides are numbered 21–23. **c-d** ABE- and PE-generated frequencies of A•T-to-G•C edits at endogenous HEK3 and FANCF sites. **e-f** Indel frequencies from the experiments in A and B. **f-g** Indel frequencies from the experiments in C and D.

### Correction of a pathogenic mutation in iPSCs with PEs and BEs

We next applied PEs and BEs to correct a disease-causing mutation in a patient-derived iPSC line carrying a homozygous PiZZ mutation (1024 G>A, E342K) in the serpin family A member 1 (*SERPINA1*) gene, the most common mutation causing α 1-antitrypsin (A1AT) deficiency^21, 22^. We first tested the ability of an ABE to correct this mutation. In this experiment, we used an NG PAM-targetable ABEmax (named NG-ABEmax) and selected a sgRNA in which the target nucleotide is the seventh adenine from the 5’ end (A7). High-throughput sequencing data from NG-ABEmax-treated iPSCs showed that this ABE yielded A-to-G conversions at a frequency of 2.5%, with a 3.7% conversion rate for a bystander adenine at the 5th position that ultimately results in an Asp-to-Gly mutation (Fig. 5a). When we considered the frequency of the intended edit alone (i.e., the A7 correction without bystander editing), it was only 0.85% (Fig. 5b).

**Fig. 5.**
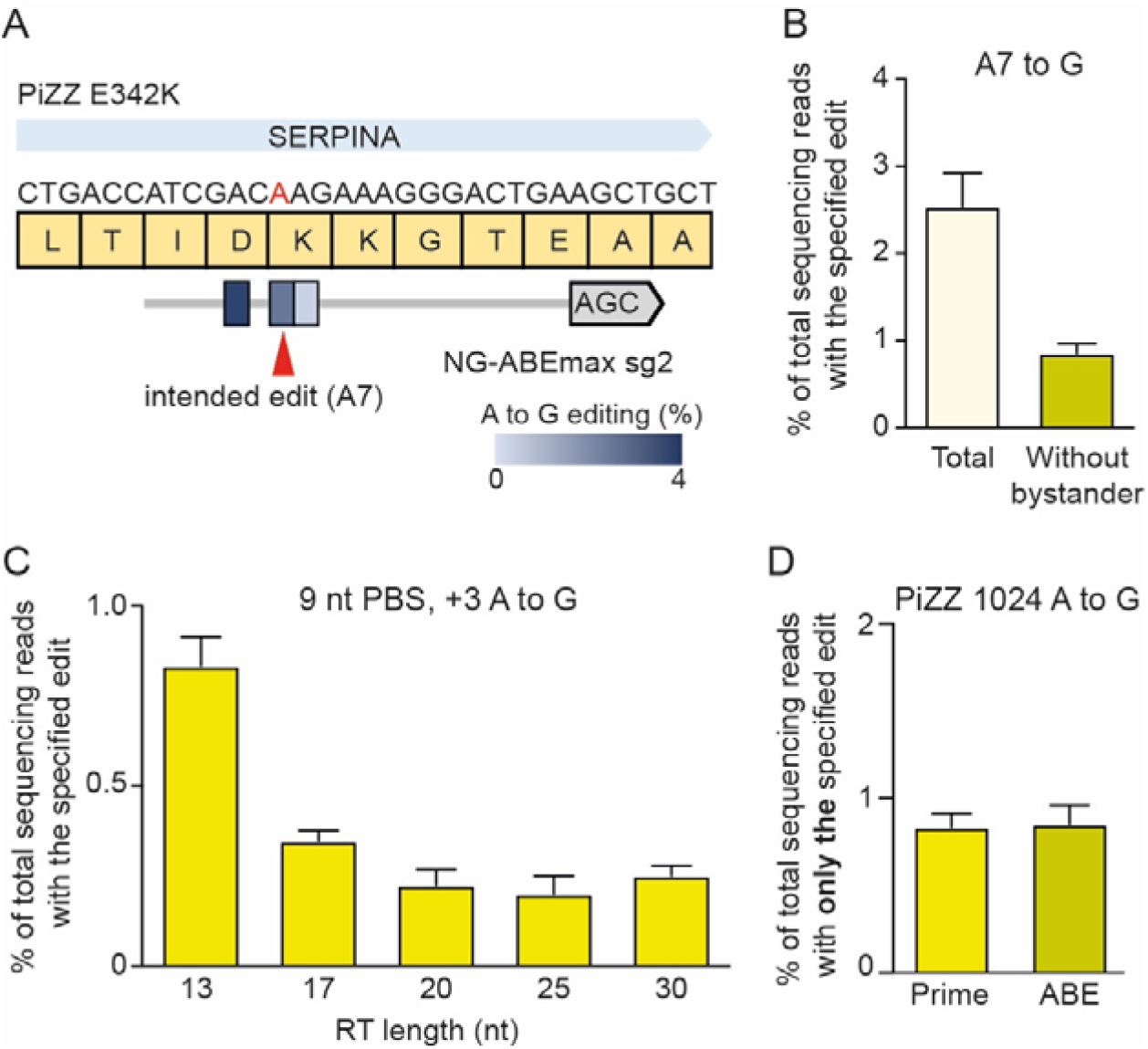
Repair of the PiZZ 1024 G>A mutation in induced pluripotent stem cells (iPSCs) derived from a patient with alpha-1-antitrypsin deficiency. **a** Schematic of the disease-associated 1024 G>A mutation in the *SERPINA1* gene, the altered protein sequence (E342K), and the sgRNA used to correct the mutation with NG-ABEmax. **b** Frequencies of A-to-G conversions in the editing window induced by NG-ABEmax. **c** Frequences of A-to-G conversions at the G>A mutation site induced by NG-PE2. pegRNAs with varying RT lengths were tested. **d** Comparison of the frequencies of precise correction of the PiZZ mutation in patient-derived iPSCs by NG-PE2 and NG-ABEmax.

We next tested a version of PE2 that recognizes the canonical PAM (NGG) for correction of the PiZZ mutation. We designed pegRNAs with varying PBS lengths, with the targeted adenine at the +24 position. We then electroporated the pegRNAs with an additional nicking sgRNA into the patient-derived iPSCs; however, we did not observe any detectable editing (Supplementary Fig. 5), similar to a previous study in liver organoids^23^. Therefore, we also tested an NG PAM-targetable PE2 (NG-PE2), which changed the edit position from +24 to +3. This PE3 system led to a low but detectable frequency of gene correction (0.83%) (Fig. 5c). Thus, in this experiment, PE and ABE demonstrated comparable efficiencies of generating only the desired edit (Fig. 5d).

### PE is a safe editing tool in hPSCs

BEs and PEs are similar in that they both contain nCas9 domains for binding target sequences, but differ in their functional components: BEs include a cytosine/adenine deaminase and PEs a RT. Previously, several groups reported that the cytosine deaminases in CBEs could generate sgRNA-independent DNA/RNA conversions, whereas the adenine deaminases in ABEs could generate sgRNA-independent RNA conversions^14-17^. Likewise, we wondered whether PEs would exhibit pegRNA-independent RT enzyme effects. To the best of our knowledge, the effects of prolonged PE expression in hPSCs have not been reported so far. Therefore, we set out to investigate whether long term PE expression would cause undesired mutations (Fig. 6a). The H9-iPE2 cells were maintained in the absence or presence of dox for 3 weeks, after which individual clones were selected and cultured in the absence of doxycycline. After expansion of the cells, genomic DNA was isolated and subjected to whole genome sequencing (WGS). We did not find any increase in the number of single nucleotide variations (SNVs) or indels in clones in which PE2 was overexpressed compared to control clones (Figs. 6b and 6c), suggesting that pegRNA-independent off-target effects in DNA might be negligible in hPSCs.

**Fig. 6.**
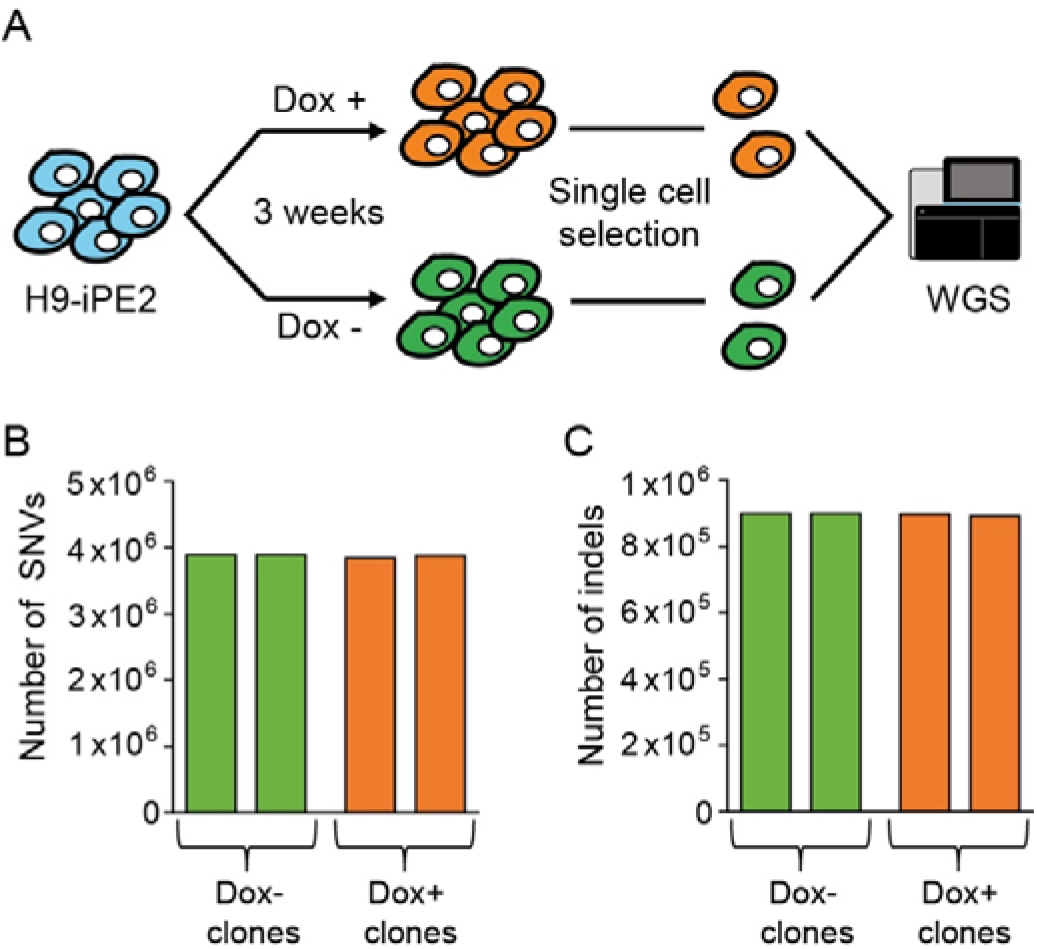
PE does not lead to genome-wide off-target effects. **a** Schematic diagram of the experimental design for clonal expansion and whole genome sequencing analysis of H9-iPE2 cells with or without dox-induced PE2 expression for 3 weeks. **b-c** Numbers of mutations in uninduced and induced H9-iPE2 clones. The numbers represent all sequence variations, including indels and single nucleotide variations.

## Discussion

Since its first demonstration in cancer cells and mouse cortical neurons, prime editing has been applied to various organisms and model systems including plants, human intestinal and liver organoids, Drosophila, mouse embryos, and adult mice^23-32^. Here, we expanded this list to include hPSCs. In this study, we demonstrated prime editing in hESCs using a dox-inducible, PE2-expressing line. Through comprehensive tests and analysis of PE3 activity at endogenous targets, we observed several strengths and weaknesses of this system in hESCs, as follows: i) PEs are capable of introducing all types of base substitutions at positions as far as 30 bps downstream of the nicking site in hPSCs and showed a trend of activity similar to that seen in other human cell lines such as HEK293T, but in general, prime editing efficiencies are lower in hPSCs than in other cell lines, ii) Compared to PEs, BEs showed higher base conversion activities but generated bystander edits, whereas PEs provided precise base conversion without bystander edits, iii) PE3-mediated base conversion is generally accompanied by the generation of indels, which are mainly caused by the combinatory activity of RT and the pegRNA, rather than by the double nicking strategy per se, and iv) WGS results revealed that long term overexpression of PE2 in hESCs did not generate significant frequencies of genome-wide SNVs or indels. Recent studies in organoid model systems showed that prime editing produces higher desired edit/indel ratios than HDR and that base editing is superior to prime editing in terms of editing efficiency, which is consistent with our conclusions.

A potential limitation of using canonical CRISPR nucleases in clinical applications is the generation of off-target mutations at sites with high homology to the intended on-target site. However, Anzalone et al. reported that PEs produce fewer off-target mutations than Cas9 nucleases and suggested that the three hybridization steps required for prime editing [i) between the pegRNA spacer and the target DNA for Cas9 binding, ii) between the pegRNA PBS and the target DNA to initiate reverse transcription, and iii) between the RT product and the target DNA during flap resolution] may reduce off-target mutagenesis. These observations were further corroborated by two studies: nDigenome-seq revealed that PEs are highly precise^33^ and WGS analysis of prime-edited clonal intestinal organoids showed that PEs exhibited high fidelity^23^. Consistent with previous studies, we revealed here with WGS analysis that long term overexpression of PE2 produced negligible off-target effects in hESCs. These results strongly support further applications of PEs for therapeutic editing in clinical studies.

Although PEs have much lower off-target editing potential than Cas9 nucleases, undesired on-target edits such as indels have been reported for PEs, especially for the PE3 system^19^. In a previous study, PE3 led to more undesired outcomes than intended edits at the target site in mouse embryos^31^; double nicking associated with the pegRNA and sgRNA in PE3 was proposed as the primary reason for the high indel rates. In contrast, based on our observations summarized in Figure 3, we suggest that the major reason is the combinatory activity of the RT and pegRNA, rather than simply the double nicking strategy. However, the detailed mechanism responsible for indel generation was not elucidated in this study. Further characterization of the repair mechanisms that resolve the flap during prime editing would potentially provide insight into the reason for the high frequencies of PE3-mediated indel formation and allow us to manipulate the repair pathway to improve the accuracy of prime editing.

In addition, despite the unique capabilities of PEs, the complexity of pegRNA design could be a potential drawback for their use. The optimal lengths of the PBS and RT template must be determined because these sequences can have substantial effects on the efficiency of prime editing. Recently, the Kim group developed computational models for predicting pegRNA efficiency and determined parameters that influence pegRNA activity^34^. However, several pegRNAs should still be tested for each specific editing task.

In summary, prime editing adds a new dimension to CRISPR-based approaches and broadens the scope of genome editing in hPSCs by complementing other methods. It is likely that an understanding of the repair mechanisms involved in prime editing and further improvement of PEs will accelerate translation of this technology into therapeutic applications.

## Acknowledgements

This research was supported by grants from the National Research Foundation of Korea (NRF) (no. 2020M3A9I4036072, no. 2020R1A6A1A06046728, no. 2021R1A2C3012908, and no. 2021M3A9H3015389) to S.B.

## Author contributions

S.B. and O.H. conceived this project; O.H. and G.H. performed the experiments; O.H. and G.-H.H. performed bioinformatics analyses; O.H. and S.B. wrote the manuscript with the approval of all other authors.

## Additional information

Supplementary Information accompanying this paper is available at http://

## Conflict of interest

The authors declare no financial or non-financial conflicts of interest.

## Methods

### hPSC culture

Cells were maintained in Essential 8 (E8) medium (Gibco A1517001) on iMatrix-511 (Matrixome, 892 021) and fed daily. ReLeSR (Stemcell Tech., 05873) was used to dissociate hPSCs into clumps for passaging, after which cells were replated in E8 containing p160-Rho-associated coiled-coil kinase (ROCK) inhibitor Y-27632 (2 μM; Stemcell Tech, 72307).

### Generation of cell lines with inducible PE2 and Cas9 expression

The plasmid for inducible PE2 expression used in this study was derived from pAAVS1-NDi-CRISPRi (addgene, 73497) by replacing the sequences encoding KRAB-dCas9-mCherry with that encoding PE2 (amplified from pCMV-PE2; addgene, 132775). To generate iPE2 and iCas9 cell lines, two million H9 hESCs were co-electroporated with the appropriate knockin vector (5 μg) and plasmids encoding AAVS1-targeting TALENs (2 μg; addgene, 59025 and 59026) using an Amaxa 4D Nucleofector system (Lonza). Serial cell dilutions were then seeded in six-well plates in E8 supplemented with Y-27632 (10 μM). After selection with the appropriate antibiotic, clones were picked, expanded, and screened by treating with dox and staining for Cas9. For genotyping, genomic DNA was extracted with a DNeasy Blood & Tissue Kit. KOD -Multi & Epi (Toyobo, KME-101) was used for junction PCR according to the manufacturer’s protocol.

### Electroporation of pegRNA or sgRNA

iPE2 hPSCs were treated with dox (1 μg/ml) for 1 or 2 days before and during electroporation. Before electroporation, accutase (Stemcell Technologies, 07920) was used to generate a suspension of single cells. 1×10^5^ cells were electropotated with 250 ng of pegRNA-encoding plasmid and 83 ng of sgRNA-encoding plasmid (for PE3) using a NEON system (ThermoFisher) at 1,050 V 30 ms (two pulses). Cells were then seeded in 48-well plates in E8 supplemented with Y-27632 (10 μM for 24 h). Electroporation of sgRNA-encoding plasmid into the iCas9 cells was performed under the same conditions. For base editing, 750 ng of ABEmax-encoding plasmid (addgene, 124163) and 250 ng of sgRNA-encoding plasmid were co-electroporated into 1×10^5^ hiPSCs.

### High-throughput DNA sequencing of genomic DNA samples

Cells were cultured for 3 days following nucleofection, after which the medium was removed and the cells were washed with 1× phosphate-buffered saline (Wellgene, LB 001-01). For genomic DNA extraction, 50 μl lysis buffer (10 mM Tris-HCl, 0.05% SDS, 25 μg/ml proteinase K) was added directly into each well of the tissue culture plate, which were then incubating at 37 °C for 1 h. Proteinase K was then inactivated by incubation at 85 °C for 15 min. For sequencing library generation, target sites were amplified directly from genomic DNA in the lysate using a KAPA HiFi HotStart PCR kit (KAPA Biosystems, KK2501) as previously described^18^. Briefly, each target was amplified using primers containing Illumina forward and reverse adapters followed by a second amplification with Illumina barcoding primers. The second-round PCR products were pooled and purified using a PCR purification kit (GeneAll, 103-150). Libraries were quantified and subjected to paired-end read sequencing using Illumina MiniSeq platform. For sequence analysis, alignment of raw reads to a reference sequence was performed using PE analyzer (http://www.rgenome.net/pe-analyzer/#!).

### Immunofluorescence

Cells were fixed with ice cold methanol and permeabilized with 0.1% Triton X-100 in phosphate-buffered saline for 30 min at room temperature. The cells were then washed thrice with wash buffer (0.03% Triton X-100 in phosphate-buffered saline). The fixed samples were blocked for 1 h with 5% bovine serum albumin (Gendepot, A0100) in wash buffer, followed by incubation for 24 h at 4 °C with anti-Cas9 (BioLegend, 844301) primary antibody diluted in wash buffer. The samples were washed with wash buffer, and then incubated with secondary antibodies at room temperature for 1.5 h. To visualize cell nuclei, the cells were counterstained with 4′,6-diamidino-2-phenylindole (DAPI) (Thermo Fisher Scientific, R37606). Cells were then viewed and photographed using a fluorescence microscope (Zeiss, axio observer z1).

### Data Availability

High-throughput sequencing and WGS data have been deposited in the NCBI Sequence Read Archive database (SRA; https://www.ncbi.nlm.nih.gov/sra) under accession numbers *****.

## Notes

### Competing Interest Statement

The authors have declared no competing interest.

