## Supplementary Figures for "Comprehensive analysis of prime editing outcomes in human embryonic stem cells"

**Table of Contents**

**Supplementary Figure 1.** Prime editing system.

**Supplementary Figure 2.** Characterization of H9-iPE2 cells.

**Supplementary Figure 3.** Comparison of prime editing efficiencies using pegRNAs with varying PBS lengths in H9-iPE2 and HEK293T cells.

**Supplementary Figure 4.** Generation and characterization of H9-iCas9.

**Supplementary Figure 5.** Targeting the PiZZ 1024 G>A mutation in patient-derived induced pluripotent stem cells with a version of PE2 recognizing an NGG PAM.

**Supplementary Figure 1.** Prime editing system. **(A)** Schematic of PE and pegRNA. (**B)** The nuclease domain (nCas9) of the PE nicks the PAM-containing strand. The liberated 3’ end binds to the PBS and RT synthesizes edited DNA using the RT template of the pegRNA. (**C)** Elongation of the 3’ end by RT generates a 3’ flap that contains the edited sequence. The 3’ flap can then be transformed to a 5’ flap through flap equilibration. Resolution of the 5’ flap by DNA repair machinery incorporates the intended edit into the DNA.

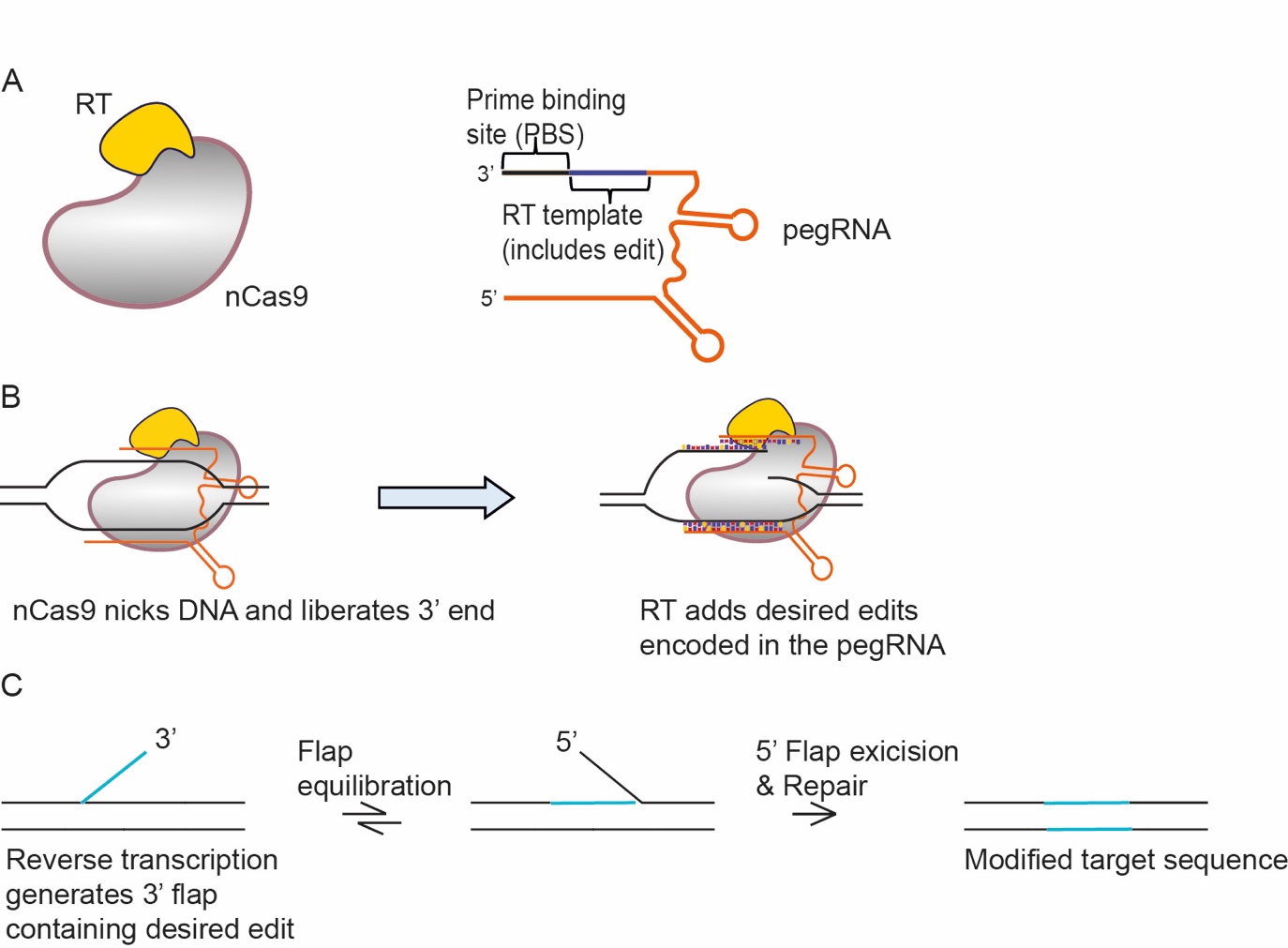

**Supplementary Figure 2.** Characterization of H9-iPE2 cells. **(A-B)** PCR-based genotyping confirmed homozygous and correct targeted integration of the inducible PE2 expression cassette into the AAVS1 locus. Non-transfected parental cells were used as a negative control for the genotyping. Colored arrows indicate the locations of the primer sets used for genotyping (top panels). The dotted rectangles delineate the areas in the gels shown in Fig. 1 (middle panels). Sanger sequencing was performed to confirm the integration (bottom panels).

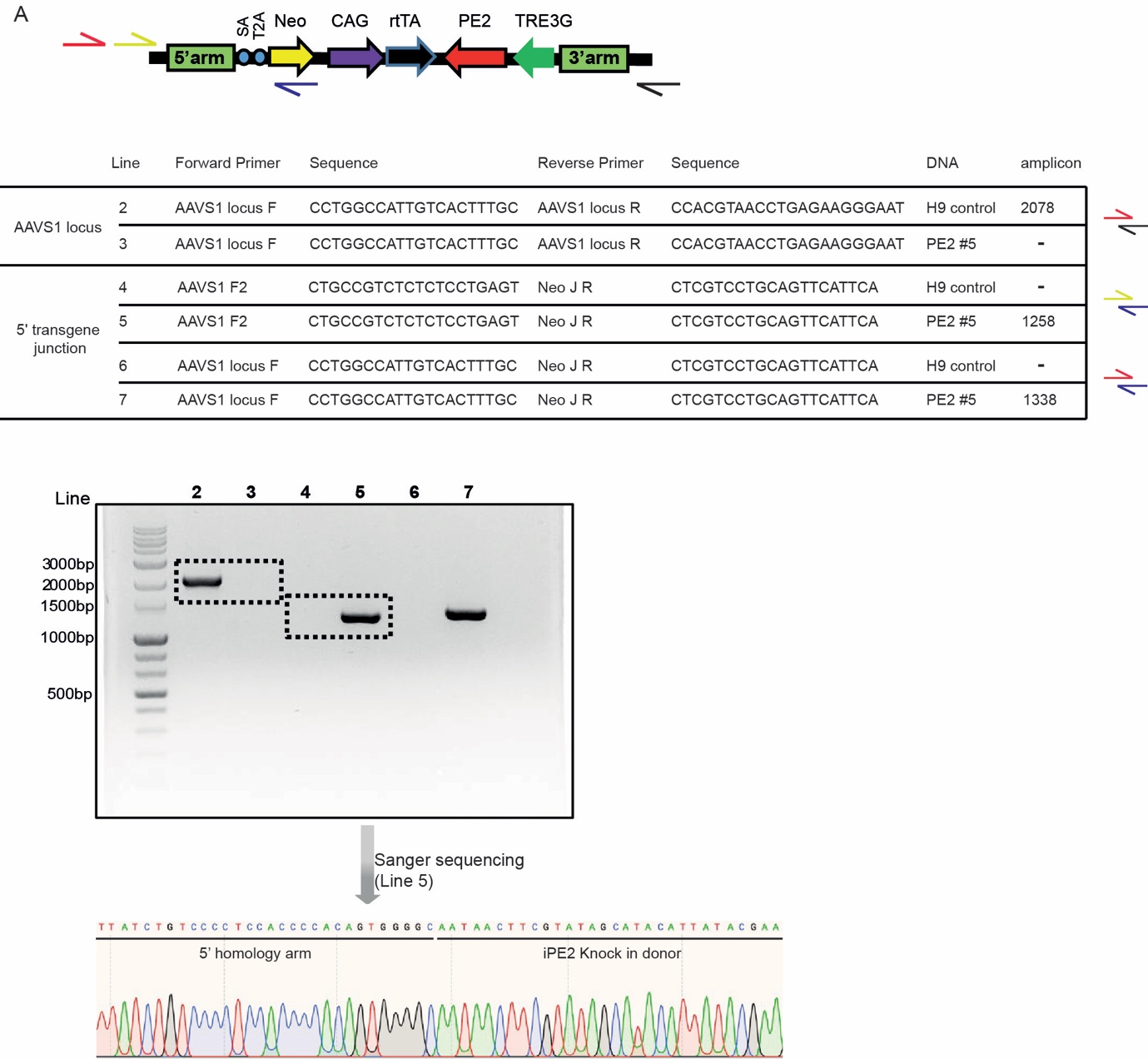

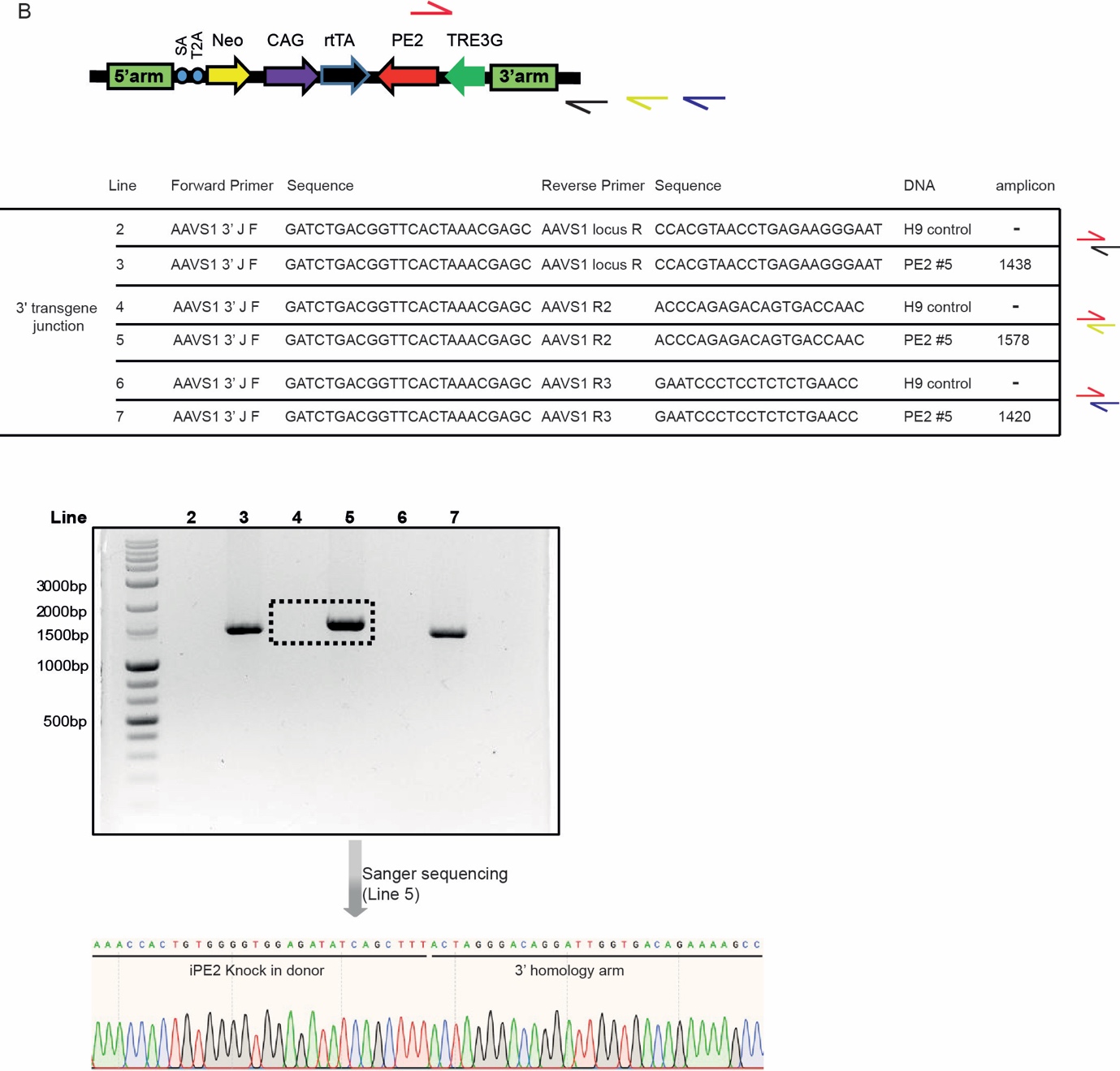

**Supplementary Figure 3.** Comparison of prime editing efficiencies using pegRNAs with varying PBS lengths in H9-iPE2 and HEK293T cells. (**A)** HEK3 site. (**B)** RNF2 site.

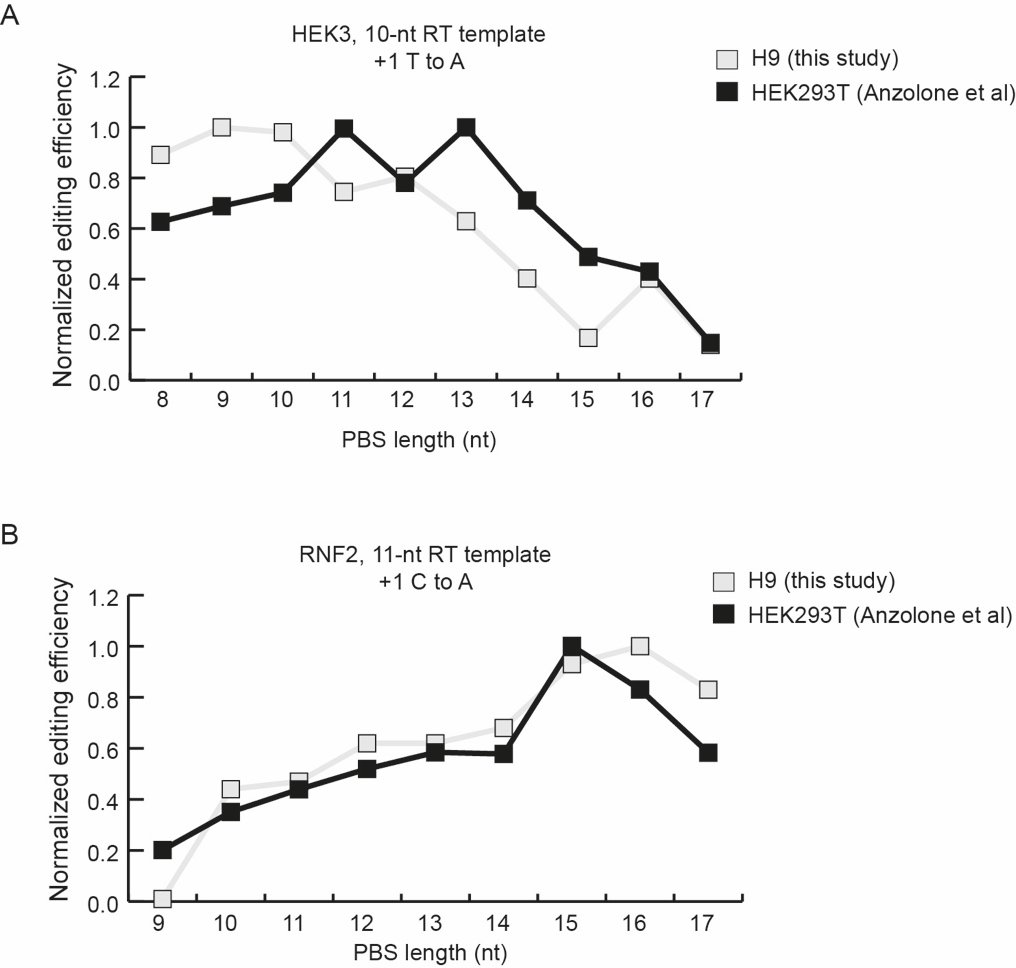

**Supplementary Figure 4.** Generation and characterization of H9-iCas9. **(A)** Schematic diagram of the strategy for TALEN-mediated targeting of the AAVS1 locus to generate H9-iCas9 cells, in which Cas9 expression is induced by dox. The AAVS1 donor vector contains a cassette in which Cas9 expression is under the control of the dox-inducible TRE3G promoter. SA, splice acceptor; 2A, self-cleaving 2A peptide; Neo, neomycin resistance gene; rtTA, dox-controlled reverse transcriptional activator; CAG, cytomegalovirus early enhancer/chicken β actin promoter. (**B)** Induction of Cas9 expression by the addition of dox. The Cas9 protein was detected by immunostaining using an anti-Cas9 antibody (green). Nuclei were stained with DAPI (blue). (**C-D)** PCR genotyping from an individual clone demonstrating successful knock-in of the inducible Cas9 expression cassette into the AAVS1 locus. Colored arrows indicate the location of the primer sets used for genotyping.

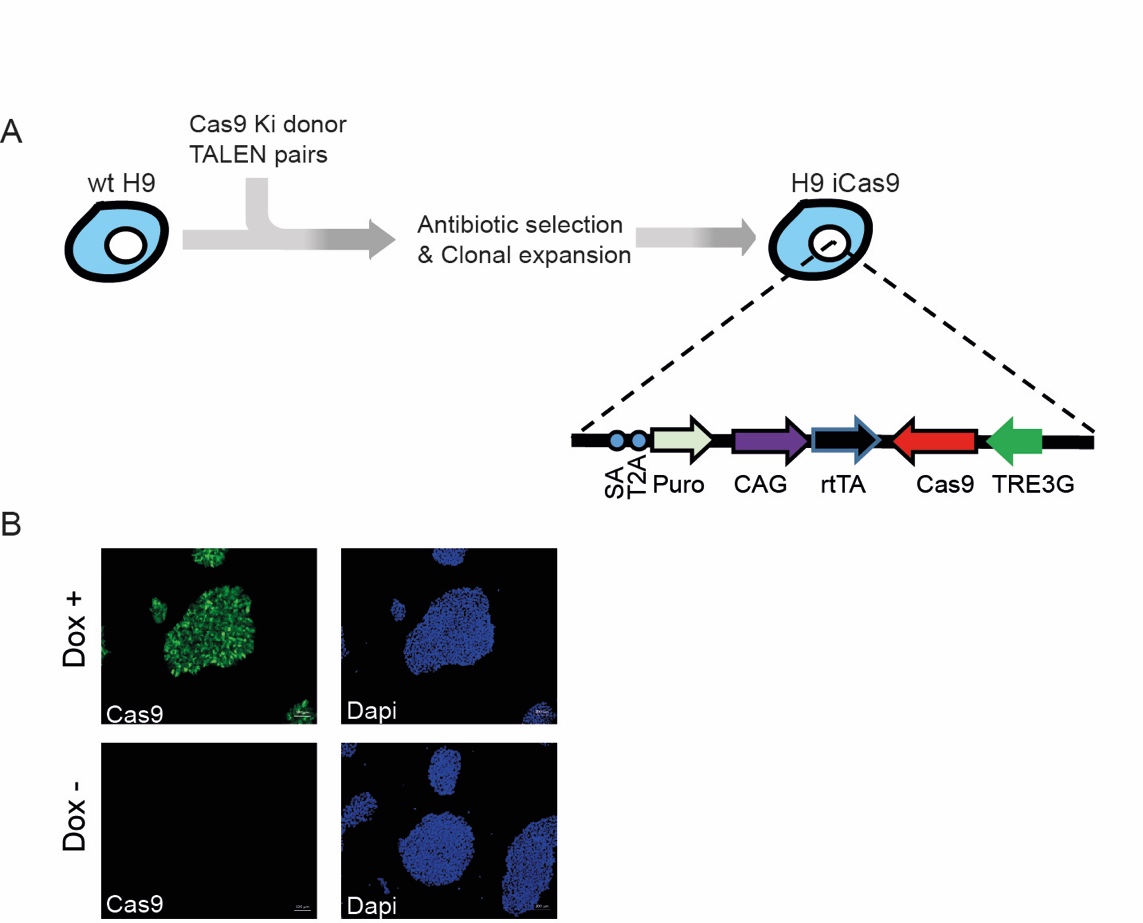

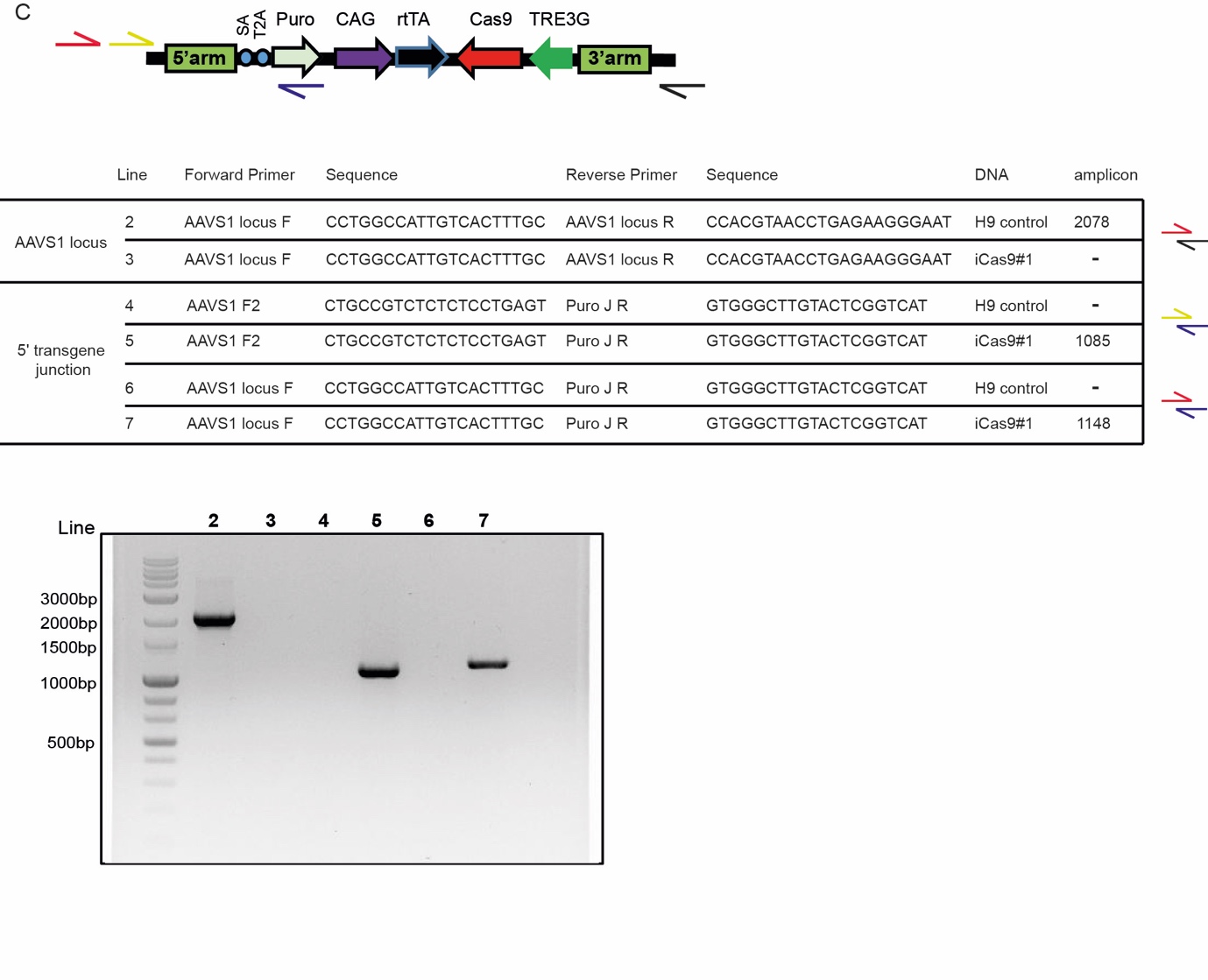

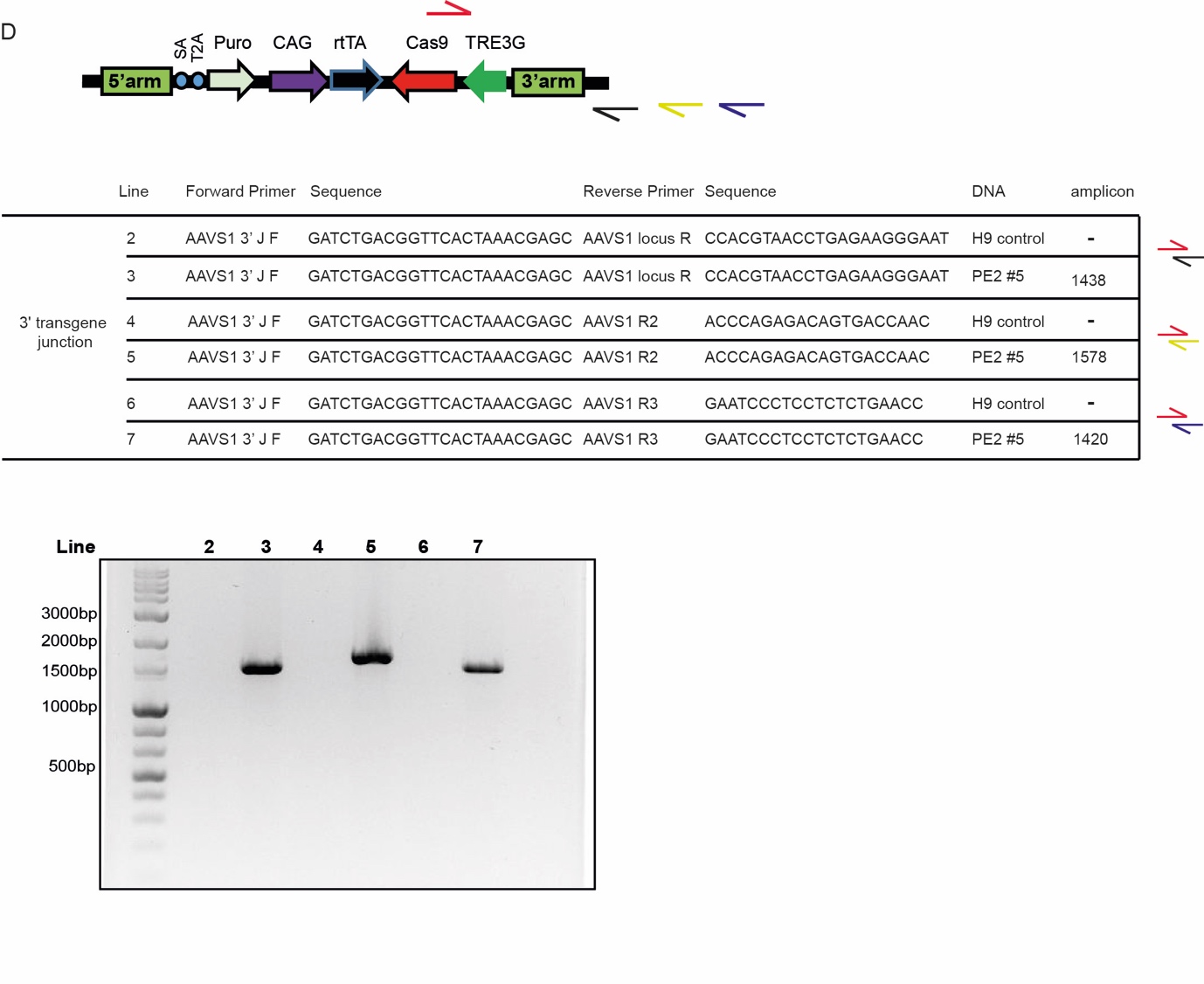

**Supplementary Figure 5.** Targeting the PiZZ 1024 G>A mutation in patient-derived induced pluripotent stem cells with a version of PE2 recognizing an NGG PAM. (**A**) The targeted adenine was at the +24 position. pegRNAs with varying PBS lengths were tested in the presence of an sgRNA, but the frequencies of the desired edit were at background levels.

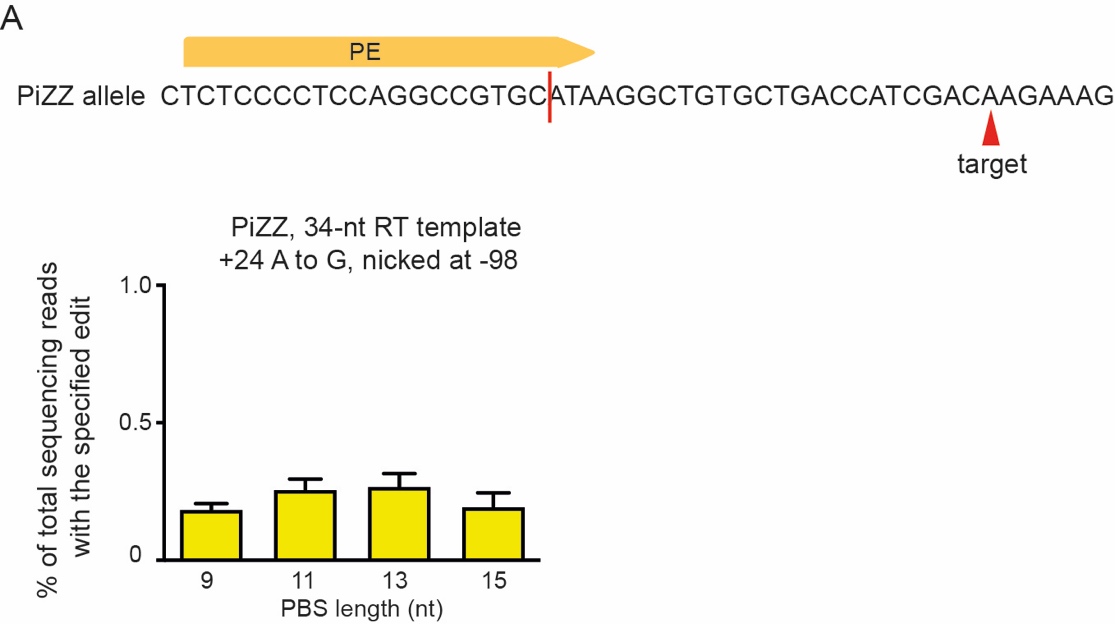
